# Effects Of Extrinsic Reward Based Skill Learning On Motor Plasticity

**DOI:** 10.1101/2024.09.10.612238

**Authors:** Goldy Yadav, Pierre Vassiliadis, Cecile Dubuc, Friedhelm C. Hummel, Gerard Derosiere, Julie Duque

**Author notes:** Conflict of interest: **None**. These authors contributed equally to this work (Goldy Yadav, Pierre Vassiliadis).

## Abstract

Human motor skill acquisition is improved by performance feedback and coupling such feedback with extrinsic reward (such as money) can enhance skill learning. However, the neurophysiology underlying such behavioral effect is unclear. To bridge this gap, we assessed the effects of reward on multiple forms of motor plasticity during skill learning. Sixty-five healthy participants divided in three groups performed a pinch-grip skill task with sensory feedback only, sensory and reinforcement feedback or both feedback coupled with an extrinsic monetary reward during skill training. To probe motor plasticity, we applied transcranial magnetic stimulation on the left primary motor cortex at rest before, during and after training in the three groups. We evaluated the amplitude and variability of corticospinal output, GABA-ergic short-intracortical inhibition and use-dependent plasticity before training and at two time points during and after training. At the behavioral level, monetary reward accelerated skill learning. In parallel, corticospinal output became less variable early on during training in the presence of extrinsic reward. Interestingly, this effect was particularly pronounced for participants who were more sensitive to reward, as evaluated in an independent questionnaire. Other measures of motor excitability remained comparable across groups. These findings highlight that a mechanism underlying the benefit of reward on motor skill learning is the fine tuning of early-training resting-state corticospinal variability.

**SIGNIFICANCE STATEMENT:** Skill acquisition is enhanced in the presence of reward. Despite its potential clinical relevance for motor rehabilitation, the underlying neurophysiological mechanisms remain largely unexplored. Specifically, whether reward affects the plasticity of motor cortex in the context of skill learning is unclear. We show that reward reduces the variability of corticospinal output at an early stage during training and that this effect correlates with individual sensitivity to reward. Our results suggest that a key mechanism underlying the beneficial effect of reward on motor skill learning may be an increase in the stability of motor output in response to training during early stages of skill learning.

## 1. INTRODUCTION

The ability to learn a wide variety of motor skills is a fundamental feature of human behavior, which is associated with plastic reorganization in the motor system (Sampaio-Baptista et al., 2018; Krakauer et al., 2019). Motor skills may emerge from distinct but inter-independent learning processes such as explicit strategies (Xeroulis et al., 2007; Wulf and Shea, 2010), implicit sensorimotor processes (Magill, 1998; Kal et al., 2018), use-dependent plasticity (Mawase et al., 2017), and reinforcement learning (Lohse et al., 2019; Vassiliadis et al., 2024). Among these, reinforcement learning, which is a well-conserved evolutionary mechanism enabling appropriate action selection based on previous outcomes (Cisek, 2019), appears to play a crucial role in guiding human motor behavior (Dhawale et al., 2017; Vassiliadis and Derosiere, 2020; Vassiliadis et al., 2021). Research indicates that providing reinforcement feedback during training can significantly improve motor skills (Abe et al., 2011; Dayan et al., 2014; Mawase et al., 2017), a finding that holds promise for clinical translation in motor rehabilitation (Widmer et al., 2022).

Reinforcement motor learning has been typically investigated by providing reinforcement feedback (*i.e.*, knowledge of performance such as whether the executed movement was a success or failure) coupled with an extrinsic reward (*i.e.*, value associated with the possible outcomes such as monetary incentives or social praise) (Wachter et al., 2009; Abe et al., 2011; Steel et al., 2016; Sporn et al., 2022). An important difference between these two aspects is that reinforcement feedback provides useful information on previous movements to guide future motor adjustments (*e.g.*, moving to a different location after failure), but coupling such feedback with an extrinsic reward does not provide any additional information that may drive learning (*e.g.*, whether a successful reach is rewarded 1 cent or 100 euros does not give any additional information on how to correct the movement per se, see Vassiliadis et al., 2021). Providing knowledge of performance improves learning by boosting intrinsic motivation to perform well (Weeks and Kordus, 1998; Thorpe and Valvano, 2002; Oppici et al., 2024) and by allowing the regulation of motor variability in response to movement outcomes (e.g., success or failure, Vassiliadis et al., 2021; Therrien et al., 2016; Wu et al., 2014; Vassiliadis et al., 2019). On the other hand, the prospect of reward can provide extrinsic motivation and improve the speed accuracy trade-off of movements, regulate aspects of motor control such as feedback control gains (Carroll et al., 2019; De-Comite 2022; Codol et al., 2023), limb stiffness (Codol et al., 2020) or movement fusion (Sporn et al., 2022). Consistent with this specific role of extrinsic reward on motor learning, recent research showed that the combination of reinforcement feedback and monetary incentives improved motor learning compared to when only reinforcement feedback was provided (Vassiliadis et al., 2021; Sporn et al., 2022). However, despite these promising findings at the behavioral level, the effect of extrinsic reward at the neural level remains largely unexplored. It is still unclear how extrinsic rewards during training modulate plasticity in the motor system.

In the present study, we investigated plasticity measures related to excitability changes in the primary motor cortex (M1) during motor skill learning. M1 is a crucial hub of the motor learning network (Reis et al., 2009; Hardwick et al., 2013; Kawai et al., 2015), which undergoes plastic reorganization in the early stages of motor learning (Pascual Leone 1995; Rioult-Pedotti et al., 1998, 2000, Classen et al., 1998; Butefisch et al., 2004; Duque et al., 2008; Vassiliadis et al., 2020). Moreover, recent research shows that M1 responds to reward (Ramakrishnan et al., 2017; Levy et al., 2020; Lee et al., 2022) and receives dopaminergic projections from the midbrain that convey reinforcement information during motor learning (Hosp et al., 2011; Leemburg et al., 2018). In addition, learning to adapt movements with knowledge of performance induces plastic changes in M1 (Uehara et al., 2018), including use-dependent plasticity (UDP) (Mawase et al., 2017) and is boosted when combined with stimulation of M1 (Spampinato et al., 2019). Finally, the presence of reward can increase corticospinal excitability (CSE) in humans (e.g., Klein et al., 2012) and modulate GABAergic short-intracortical inhibition (SICI) in M1 (Thabit et al., 2011; Spampinato et al., 2019; Hamel et al., 2023).

Based on these findings linking M1 and reward, we set out to evaluate skill training related plasticity in M1 in a subset of individuals (n=65) who participated in Vassiliadis et al. (2021) and learned a motor skill task in the presence or absence of reward. We measured amplitude and variability of CSE, SICI and UDP at different time points during the experiment with a specific focus on an early stage of training – *i.e.*, when groups are exposed to different feedback, with only one group receiving both reinforcement feedback and monetary reward – and on the end of training – *i.e.*, when all groups receive the same reinforcement feedback on the skill task with no monetary reward.

## 2. MATERIALS AND METHODS

Sixty-five right-handed healthy subjects successfully completed this study (44 Females; 23.85 ± 3.22 years old) which involved motor skill learning and neurophysiological measurements using transcranial magnetic stimulation (TMS) over primary motor cortex (M1). As mentioned earlier in the Introduction, the data of these subjects (n=65) were from a larger pool of subjects (n=90) previously exploited in a separate study where we specifically studied behavioral changes underlying motor skill learning associated with monetary reward (Vassiliadis et al., 2021). In this current paper, we focus on motor plasticity measured in the form of motor excitability changes accompanying this form of reward related skill learning. Subjects filled out a TMS safety questionnaire to look for any contraindication and gave written informed consent in accordance with the Ethics Committee of the University (approval number: 2018/22MAI/219) and the principles of the Declaration of Helsinki. Handedness of these subjects was assessed via Edinburgh Handedness inventory (Oldfield 1971). In addition, all subjects completed a French adaptation of a short version of the Sensitivity to Punishment and Sensitivity to Reward Questionnaire-SPSRQ (Torrubia, 2001; Lardi et al., 2008) in the beginning of the experiment. None of the subjects suffered from any neurological disorder or had any history of psychiatric illness, drug, or alcohol abuse, and none reported undergoing any drug treatment that could bias their performance or their underlying neural activity. Individuals had normal or corrected vision. All subjects were financially compensated and naïve to the purpose of the study.

### 2.1 Experimental Design

#### 2.1.1 Motor skill learning Task Task Apparatus

Subjects were seated in a quiet and dimly lit room, approximately 60 cm in front of a cathode-ray tube computer screen (100 Hz refresh rate). The latter was used to display the motor skill learning task implemented using Matlab 7.5 (the Mathworks, Natick, Massachusetts, USA) and Psychophysics Toolbox extensions (Brainard, 1997; Pelli, 1997). Subjects were seated with their forearms positioned in pro-supination on a table placed in front of them. With the arms in this position, they were able to pinch a manipulandum (Arsalis, Belgium) with the thumb and index fingers, as required by the task. Subjects were explicitly asked to keep their eyes open, with the gaze oriented towards the screen (not the manipulandum), throughout the entire experiment.

##### Task Design

The motor skill task consisted of a previously described force modulation paradigm (Vassiliadis et al., 2021; Vassiliadis et al., 2022). Briefly, this task required participants to squeeze the force manipulandum using the thumb and index fingers to control a cursor displayed on the screen. Increasing the force resulted in the cursor moving vertically upward (see Figure 1A). Each trial started with a *preparatory period* in which a fixed target (7 cm diameter) appeared at the top of the screen and a sidebar (9 x 15 cm) appeared at the bottom. After a variable time interval (0.8 to 1 s), a black cursor (1 cm diameter) became visible in the sidebar, indicating the start of the *movement period*. Subjects had to pinch the manipulandum in order to move the cursor as quickly as possible from the sidebar to the target and maintain it there for the rest of the movement period lasting for 2 s. The level of force required to reach the target (Target_FORCE_) was individualized for each participant estimated at the start of the experiment and set at 10% of maximum voluntary contraction (MVC). Notably, squeezing the manipulandum before the appearance of the cursor was considered as an anticipation and led to interruption of the trial. Anticipation trials were rare given the variable preparatory period and were discarded from further analyses. On 90 % of trials, the cursor disappeared shortly after the start of the movement period when the generated force became larger than half of the Target_FORCE_ (*i.e.,* 5 % of MVC)). Hence, subjects had to learn to approximate the Target_FORCE_ in the absence of full visual feedback in order to perform appropriately in these partial vision trials. In the remaining 10% of trials, the cursor did not disappear (full vision trials, not included in analysis). Partial visual feedback was used here in order to increase the impact of other forms of performance feedback (detailed later) on learning (Izawa et al., 2011, Mawase et al., 2017, Vassiliadis et al., 2021).

**Figure 1.**
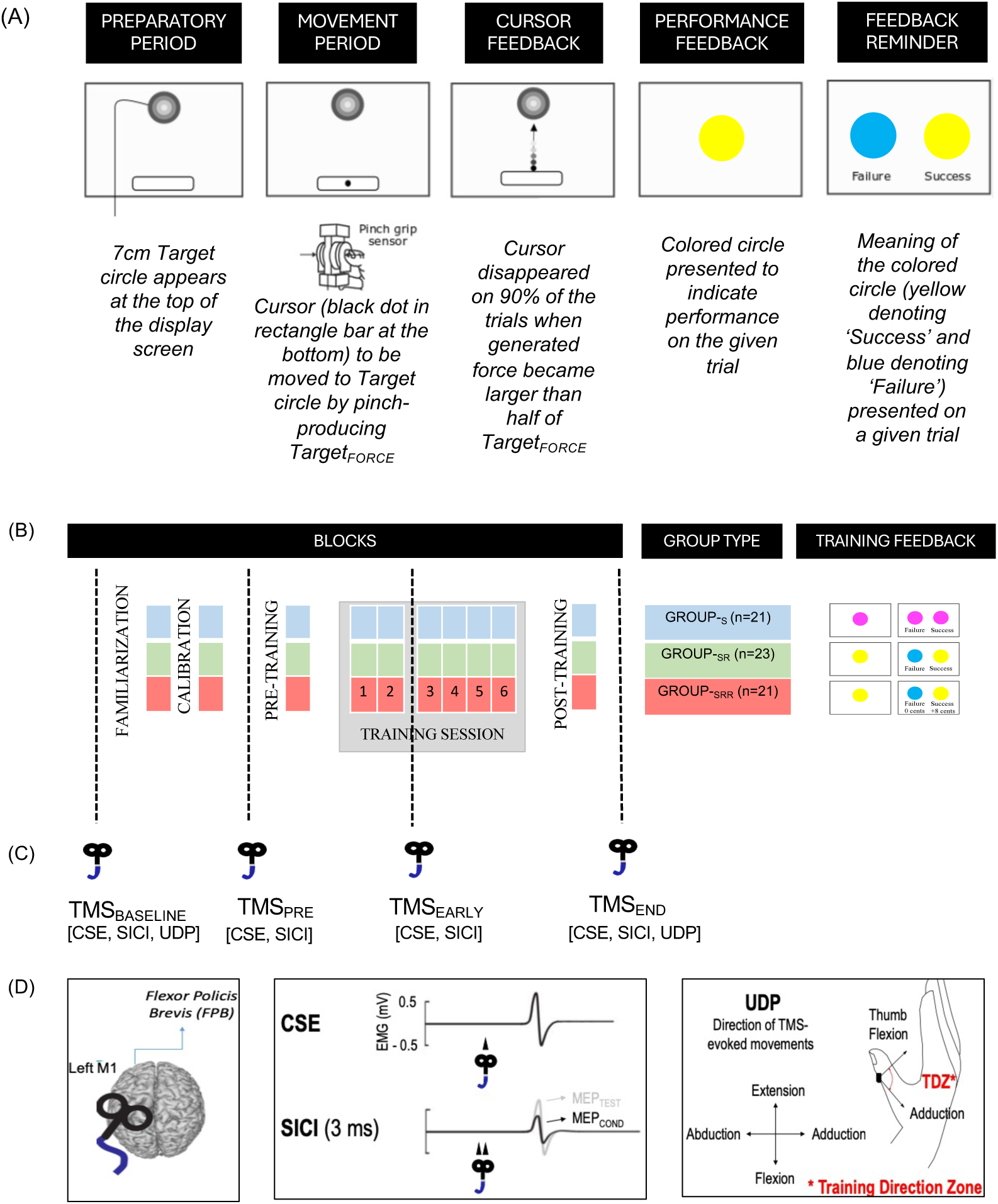
**(A)** Motor skill task depicting a target circle (top) and a rectangle start position (bottom). Participants were required to move the cursor (black dot) from the start position to the target circle by pinching the transducer and producing Target_FORCE_. At the end, performance feedback was provided in the form of colored circles (yellow or blue) with a reminder screen presenting the meaning of the colored circle (blue denoting ‘failure’ and yellow denoting ‘success’). (B) Study design depicting all the blocks for the skill task (Familiarization, Calibration, Pre-Training, Training Session and Post-Training). All three groups followed the same task design except that during the Training Session they received different performance feedback depending on the group type-Group-_S_ (in blue) received only sensory feedback, Group-_SR_ (in green) received sensory and reinforcement feedback, and Group-_SRR_ (in red) received sensory-reinforcement feedback coupled with monetary reward. Note that except the Training Session, all three groups received the same performance feedback (sensory-reinforcement) on all other blocks. (C) TMS measurements (CSE, SICI, UDP) obtained at four different time points-TMS_BASELINE_, TMS_PRE_, TMS_EARLY_, TMS_END_. (D) TMS was applied on the left M1 to obtain MEPs from Flexor Policis Brevis (FPB) muscle to assess CSE (MEP amplitude and variability), SICI (ratio of MEP_CONDITIONED_ and MEP_TEST_) and UDP (TMS evoked movements elicited in the training direction zone-TDZ, i.e., thumb adduction and flexion in our motor skill task).

To evaluate task performance, we calculated an “Error” parameter for each trial. The Error was defined as the mean of the difference between the exerted force and the Target_FORCE_ between 0.15 s and 2 s after trial onset and was expressed in %MVC. Hence, the more the force differed from the Target_FORCE_, the higher the Error on a given trial. The first 0.15 s were not considered for computing the Error because we assumed that this was the minimum time required for subjects to react to the appearance of the cursor (see Steel et al., 2016). A trial was then classified as *‘Success’* or ‘*Failure’* if the Error was under an individualized success threshold (determined during *Familiarization* session, detailed next) to provide *performance feedback* in the form of colored circles (along with a *feedback reminder* screen). Importantly, subjects were explicitly told that task success depended on their ability to approximate the Target_FORCE_ as fast and accurately as possible. In summary, for successful task performance (i.e. to have a low Error on this skill task), subjects had to quickly initiate the force and be as accurate as possible in reproducing the Target_FORCE_ to move the cursor to target and maintain it there for 2 s.

##### Task Blocks and Groups

Subjects first performed a *Familiarization* block which consisted of 20 full vision trials (not used for analysis) to allow the subjects to become acquainted with the task. After this block, the remaining blocks consisted of 90% partial vision trials that were considered for analysis. Once familiarized, subjects performed a *Calibration* block of 20 trials, which served to individualize the difficulty of the task for each subject for the rest of the experiment. As such, for each individual subject, partial vision trials of the *Calibration* block were sorted in terms of Error from the lowest to the greatest in percentage of MVC. We took the 35^th^ percentile of the Error to determine the individual success threshold (see Vassiliadis et al 2021 for details on calibration procedure). For the rest of the experiment, trials with an Error lower than this threshold were considered as ‘*Success*’ while trials with higher Error values than this threshold were considered as ‘*Failure*’. Following *Calibration*, subjects performed a total of 280 trials (8 blocks) of the skill task, out of which 20 trials in the beginning and 20 trials at the end served to assess performance on *Pre-Training* and *Post-Training* blocks, respectively. At the end of each trial of these blocks (i.e., *Familiarization, Calibration, Pre-Training and Post-Training*, see Figure 1B), subjects were presented with knowledge of performance feedback - a yellow or blue colored circle (presented for 1 s) immediately followed by a reminder screen (presented for 1.5 s) with both colored circles indicating *‘Success’* and *‘Failure’* respectively on a given trial (Figure 1A). In between the *Pre-Training* and *Post-Training* blocks, there were six training blocks (blocks 1-6) of 40 trials each. During this 6-block training session, subjects were divided into three groups namely, Group-_S_ (n=21), Group-_SR_ (n=23) and Group-_SRR_ (n=21), depending on the performance feedback they received (Figure 1B). Individuals in Group_-S_ received only somatosensory feedback to perform the task during this session: participants were explicitly aware that performance feedback was non-informative (magenta circles were displayed regardless of the performance). Group_-SR_ continued receiving the knowledge of performance feedback in the form of yellow or blue circles (indicating *‘Success’* or *‘Failure’*). For Group_-SRR_, this knowledge of performance was coupled with a monetary reward-8 cents and 0 cent for ‘*Success’* and ‘*Failure’* respectively for each trial. Therefore, contrary to Group_-S_, Group_-SR_ and Group_-SRR_ continued receiving knowledge of performance feedback; with this feedback coupled with a monetary reward only in Group_-SRR_ during the six blocks of training. However, note again that during *Familiarization*, *Calibration, Pre-Training* and *Post-Training* blocks, irrespective of the group type, all subjects received both sensory and reinforcement feedback for their performance.

The six training blocks were separated by breaks of 1.5 min to prevent fatigue and by a 7-min TMS session to probe motor plasticity at the end of training block-2 (detailed in the next section). Finally, subjects received a fixed show-up fee corresponding to 10 euros/hour of experiment. In addition, participants also gained a monetary bonus-set at 10 euros for subjects in Group_-SR_ and Group_-S_, while it was variable from 0 to 20 euros according to performance for the Group_-SRR_ (gain of 8 cents per successful trial in training blocks 1-6). Importantly, this bonus for Group_-SRR_ was determined to match that obtained by the other two groups and finally it corresponded to 9.20 ± 3.29 Euros. A t-test revealed that the total remuneration at the end was not different across the three groups (t_(21)_ = −1.11; p = 0.28).

#### 2.1.2 Transcranial magnetic stimulation (TMS) TMS Procedure

As mentioned in the introduction, our main objective was to assess M1-related neuroplasticity changes underlying reward-based skill learning. To do so, TMS pulses were delivered at four different resting time points during the experiment (Figure 1C), eliciting MEPs (1) before task *Familiarization* (TMS_BASELINE_), (2) immediately before *Pre-Training* (TMS_PRE_), (3) early during training session at the end of block-2 (TMS_EARLY_), and (4) after *Post-Training* block (TMS_END_). Subjects were asked to remain relaxed and motionless with their feet flat on the ground and the eyes open during TMS assessments. TMS was delivered over left M1 through a 70 mm figure-of-eight coil connected to a Magnetic Stimulator Bistim^2^ (i.e. two Magnetic Stimulator 200^2^ combined through a connecting module, allowing also to deliver paired pulses through one coil; Magstim, Whitland, Dyfield, UK). The coil was placed tangentially on the scalp with the handle oriented toward the back of the head and laterally at a 45-degree angle away from the midline, approximately perpendicular to the central sulcus. After fitting the participant with a headcap (Electro-cap, Electro-Cap International, USA), the left M1 “hot spot” was identified by searching the optimal scalp position at which a single pulse of TMS consistently produced detectable motor-evoked potential (MEP) in the contralateral (right) Flexor Pollicis Brevis (FPB), an agonist muscle required in our motor skill learning task. We marked this location on the cap to provide a reference mark throughout the experiment (Vassiliadis et al., 2020; Derosiere et al., 2022; Wilhelm et al., 2024; Neige et al., 2023), and proceeded to determine the resting Motor Threshold (rMT) for each subject, which corresponds to the minimal TMS intensity required to evoke MEPs of ∼50 µV peak-to-peak in the target muscle (*i.e.*, right FPB) in at least 5 out of 10 consecutive stimulations (Grandjean et al., 2018; Vassiliadis et al., 2018). Further, TMS intensity required to achieve MEPs of 1 mV in FPB was evaluated. Finally, a three-dimensional accelerometer (Kistler Instrument, Amherst, NY) was fixed on the distal interphalangeal joint of the thumb to determine the direction and amplitude of each TMS-evoked movement to assess use-dependent plasticity at the end of training (UDP, detailed in next Section).

To assess skill learning related motor plasticity, we applied single-pulse TMS on the M1 hotspot for the FPB muscle at an intensity of 130% of the rMT. Single-pulse stimulations were delivered to obtain MEPs at the four time points-TMS_BASELINE_ (60 pulses), TMS_PRE_ (20 pulses), TMS_EARLY_ (20 pulses) and TMS_END_ (60 pulses). These MEPs were used to assess changes in corticospinal excitability (all time points) as well as UDP (TMS_BASELINE_ and TMS_END_ time points). The time interval between stimulations was random and ranged from 4.5 to 5.5 seconds to prevent subjects from anticipating the pulses. Next, we wanted to assess the activity of M1 intracortical circuits throughout skill learning in the different groups of subjects. To do so, we used a paired-pulse TMS protocol on M1-a subthreshold conditioning stimulus (at 80 % of the rMT), followed by a suprathreshold test stimulus (at Intensity1mV). With this protocol, response of the test stimulus can be modified by the conditioning stimulus depending on the time interval between the two stimuli- a decrease in the test stimulus amplitude is generally observed between 2 and 6 ms, which is believed to reflect the activation of the GABAergic inhibitory circuits within M1 and the phenomenon, known as Short-latency Intracortical Inhibition (SICI) (Derosiere and Duque, 2020). In this experiment, 15 single-pulses (at Intensity1mV) and 15 paired-pulses (conditioning pulse at 80%rMT and test pulse at Intensity1mV with an interval of 3ms) were delivered in a random order to prevent anticipation and we measured the ratio between MEPs obtained from conditioned (test pulse during paired-pulses trials) and unconditioned (test pulse during single-pulse trials) stimuli (MEP_CONDITIONED_/MEP_TEST_). Time interval between the two subsequent stimuli randomly varied between 4.5 and 5.5 seconds. These measurements were also obtained at four different time points, i.e., TMS_BASELINE_, TMS_PRE_, TMS_EARLY_ and TMS_END_.

##### Electromyography (EMG) Recording

EMG was recorded to measure peak-to-peak amplitude of MEPs using surface electrodes (Ambu BlueSensor NF-50-K/12/EU, Neuroline, Medicotest, Oelstykke, Denmark) placed over right FPB, with one electrode placed on the body of the muscle and another on the distal interphalangeal joint; the ground electrode was placed on the styloid process of the ulna. The raw EMG signal was amplified (gain of 1K), band-pass filtered on-line (10-500 Hz, Neurolog; Digitimer), notch-filtered (50 Hz) (Digitimer D360, Hertfordshire, UK) and digitized at a sampling rate of 2 kHz (CED 1401-3 ADC12 and Signal 6 software, Cambridge Electronic Design Ltd) for off-line analysis. Data extraction was performed using Signal 6 and Matlab 2018a (Mathworks, Natick, Massachusetts, USA), respectively.

### 2.2 Statistical Analysis and Endpoint Measures of Interest

For behavioral performance, we assessed motor skill learning in the 65 subjects who received TMS over M1 (for the original behavioral data set of 91 subjects, see Vassiliadis et al., 2021), and compared skill learning across the three groups of subjects (group-type: Group_-S_, Group_-SR_, Group_-SRR_). Next, related to the goals of this paper we assessed three TMS-related markers of motor plasticity-MEP amplitudes (mean and variability), SICI and UDP (Figure 1D). To perform group comparisons, learning-related changes in these measures of interest were separately assessed for two main time points critical to our research objectives, i.e., end-training (TMS_END_) and at an early-training time point during the training session (TMS_EARLY_). As mentioned in the Introduction, TMS_EARLY_ allowed us to examine motor excitability measures when the three groups are learning the skill task with different conditions during the training session, while TMS_END_ allowed to examine these measures when all groups are provided the same knowledge of performance feedback (but no reward). Assessing data from these two time points thus enabled us to probe the underlying neurophysiological states depending on the task conditions. For analysis, data obtained for TMS_END_ and TMS_EARLY_ were normalized and expressed in percentage (%) of TMS_PRE_ (except in case of UDP measure for which TMS-evoked movements at TMS_END_ were directly compared with TMS_BASELINE_, see details in 2.2.4) for performing group comparisons. Additionally, with respect to Group_-SRR_ which received reward during the 6-block training session, we computed the sensitivity to punishment and reward score obtained from each subject at the beginning of the experiment (based on the SPSRQ) to perform further correlations.

Statistical analyses were performed using JMP software. Analysis of variance (ANOVA) was the primary statistical test for comparing various measures (behavior and neurophysiology) obtained for the three groups. Significance level was set at 0.05, and a significant effect on the ANOVA was further assessed by performing Tukey’s post-hoc tests. Effect sizes were reported using partial eta-squared (η^2^_p_) measure.

#### 2.2.1 Motor skill learning

We evaluated skill learning by computing the rate at which Error decreased over training blocks 1 to 6. To compute this value for each individual, we performed a linear fit of the Error data using equation: SkillError = k(BlockNumber) + C, where k is slope/rate and C is intercept. We obtained the slope (k) and intercept (C) values for each subject and compared those for the three groups by performing ANOVA. In addition to the rate of skill learning, we also computed and compared the skill Error at *Post-Training* block (in % of *Pre-Training*) for the three groups using a one-way ANOVA as done previously (Vassiliadis et al., 2021).

#### 2.2.2 MEP Amplitude (mean and variability)

To assess CSE-changes during training, we extracted the peak-to-peak MEP amplitude as well as the Root Mean Square (RMS) of EMG activity in the 200 ms preceding TMS pulse. We removed the first MEP of each block, as well as any MEP preceded by a significant muscular activity (based on RMS, threshold set at RMS > 0,02 mV). A total of 0.18% of the MEP trials were removed based on this criterion. Next, we removed outlying trials (± 2.5 SD of the mean of each block), corresponding to 2.02% of all trials. After removing these outliers, 97.8% of the MEP data set remained with at least 18 MEPs for each TMS time points for each subject. Next, we performed two analyses on this data set: we calculated the mean MEP amplitude (MEP_MEAN_) as well as the variability of corticospinal output by computing the coefficient of variation (MEP_CV_ calculated as SD/ MEP_MEAN_) (based on Vassiliadis et al., 2018; Klein Flugge et al., 2013). As our goal was to better understand excitability-related changes at two main time points with distinct behavioral requirements in our experiment, we performed one-way ANOVA for group comparisons on MEP_MEAN_ and MEP_CV_ measures, separately for TMS_END_ and TMS_EARLY_ time points (both in %TMS_PRE_). Finally, we evaluated the association between training-related changes in corticospinal excitability variability (MEP_CV_, see results) and sensitivity to reward and punishment scores with specific focus on the reward group (Group_-SRR_) by performing Pearson bivariate correlations.

#### 2.2.3 Short intracortical inhibition (SICI)

As described earlier, to evaluate SICI we alternated between 15 single-pulse (MEP_TEST_) and 15 paired-pulse (MEP_CONDITIONED_) stimuli. We pre-processed the data the same way as for the single pulse MEPs leading to the removal of 1.06% of MEP_TEST_ and 0.03% of the MEP_CONDITIONED_ data. Next, we computed SICI by obtaining the ratio (MEP_CONDITIONED_/MEP_TEST_). In this way, smaller value (less than 1) indicates intracortical inhibition and a value of 1 or higher indicates no inhibition at all and even facilitation. We had to remove seven subjects who had a SICI ratio > 1 at TMS_BASELINE_ (meaning the absence of SICI effect) and four participants who did not receive this type of stimulation due to technical issues. We performed further analysis on the remaining 54 subjects. Like MEP amplitudes data described above, we assessed learning-related changes in SICI ratios at two main time points-TMS_END_ and TMS_EARLY_ (both in %TMS_PRE_) and performed group comparisons with one-way ANOVA. Thus finally, a value less than 100% will indicate more intracortical inhibition whereas value equal to or higher than 100% will indicate less inhibition or even facilitation compared to pre-training SICI values.

#### 2.2.4 Use-dependent plasticity (UDP)

UDP manifests as directional changes in TMS-evoked movements that become increasingly similar to those required to execute the task during training (i.e., reaching the Training Direction Zone (TDZ), Mawase and al., 2017). As mentioned above, UDP was evaluated by means of a three-dimensional accelerometer fixed on the thumb, allowing us to extract the direction and amplitude of each TMS-evoked movement and to calculate the first peak acceleration vector in both the horizontal and vertical axis (respectively corresponding to abduction/adduction and flexion/extension), as done in previous studies (Classen et al., 1998; Duque et al., 2008; Galea and Celnik, 2009; Mawase et al., 2017). As our skill task involved pinching of a force sensor, TDZ was defined as a combination of adduction and flexion of the thumb (Figure 1D). Notably, while UDP has been mainly evaluated in the context of ballistic movements (Classen et al., 1998, Duque et al., 2008), recent work suggests that UDP can also be observed following isometric force modulation training (Mawase et al., 2017). To assess training-related plasticity changes, UDP measures were obtained at two time points: at the beginning (TMS_BASELINE_) and end of the experiment (TMS_END_). We analyzed UDP related to training on our skill task in the three groups by comparing the percentage of the TMS-evoked movements that fell into TDZ at TMS_END_ compared to TMS_BASELINE_ by performing a two-way ANOVA (with time-point and group as factors). Note that five subjects (out of 65) received the 60 pulses at TMS_PRE_, instead of TMS_BASELINE_. Notably, removing these individuals from the analysis did not change the results. All 65 subjects were therefore included.

## 3. RESULTS

BEHAVIOR

### 3.1 Rate of motor skill learning was faster when performance was coupled with reward during the training session

We found that skill learning during the training session in the three groups depended on the presence of extrinsic reward during the training session (see Error over Block 1-6 in Figure 2A). To quantify this learning, we computed the intercepts and slopes corresponding to linear fits of these Error data over the six blocks (normalized to Pre-Training block). The intercept values (Figure 2B) were comparable across all three groups [Group_-S_: 108.07±6.16, Group_-SR_: 99.13±5.88, Group_-SRR_: 110.04±6.16] as indicated by no significant group effect (F_2,62_=0.9428, p=0.3950, η^2^_p_=0.0295). Hence, all subjects started with a comparable level of performance in the task. Then interestingly, we found a negative slope only for the group that trained with reward [Group_-S_: 0.69±1.17, Group_-SR_: 1.01±1.12, Group_-SRR_: −4.17±1.17] indicating substantial reduction in Error or faster learning rate in Group_-SRR_ (see Figure 2C). Statistically, we found a significant effect of group (F_2,62_=6.2759, p=0.0033, η^2^_p_=0.1683) on the slope. Tukey’s post-hoc tests revealed that the slope for Group_-SRR_ was significantly higher compared to Group_-SR_ (p=0.0060) as well as Group_-S_ (p=0.0126), while there was no significant difference between Group_-SR_ and Group_-S_ (p=0.9789).

**Figure 2.**
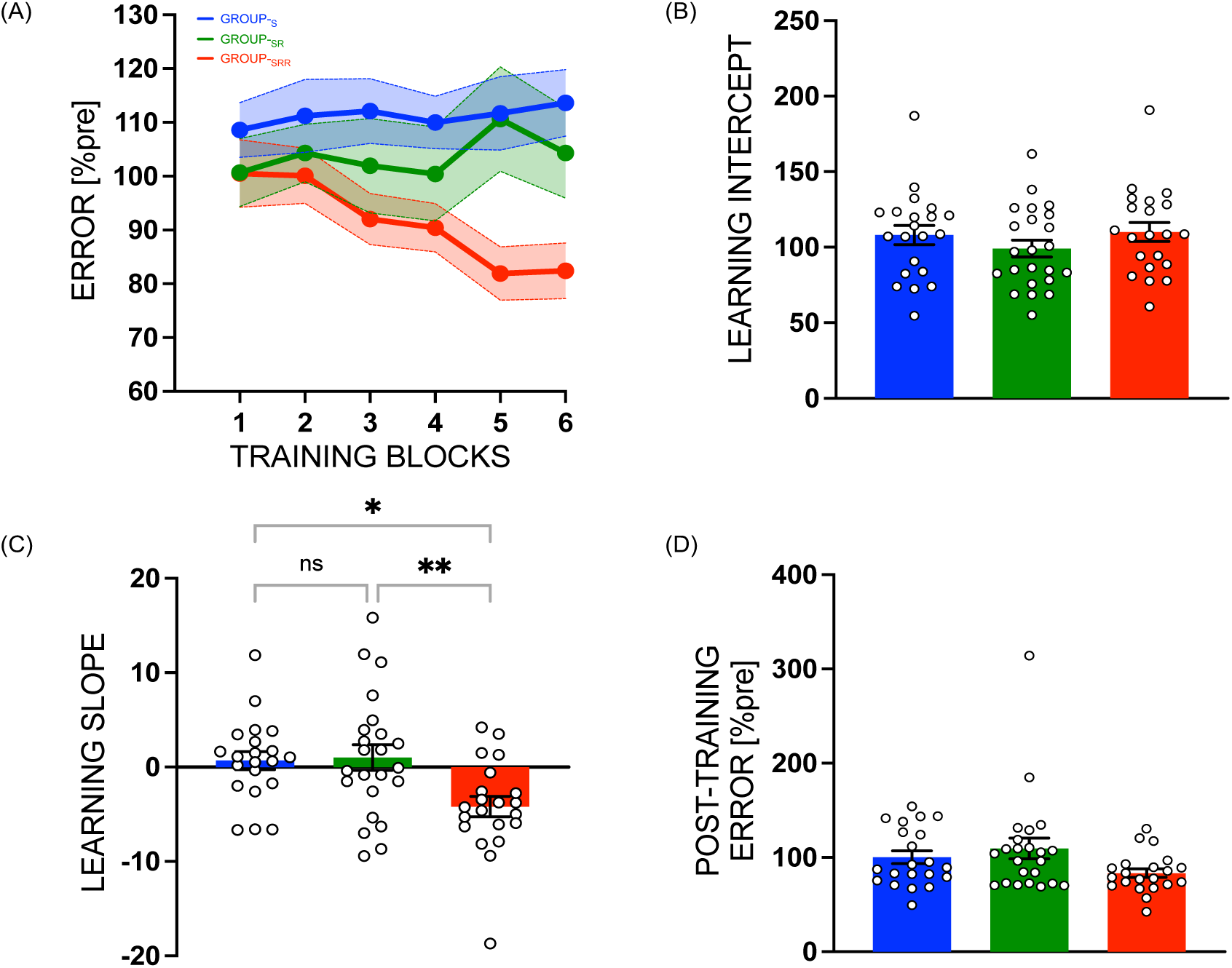
Faster motor skill learning in presence of reward. (A) Motor skill performance of the three groups (Group-_S,_ Group-_SR,_ Group-_SRR_) during six blocks of training session, with (B) no significant difference in learning intercept. (C) Rate of learning was significantly different in the group that trained with reward (Group-_SRR_) as compared to the other two groups. (D) Post-training error (when all groups perform the task with same performance feedback) was not significantly different across groups. *indicates p<0.05, **indicates 0.005<p<0.05. Data represent mean, normalized to Pre-Training block. Error bars/bands represent Standard Error. White circles (with black outline) represent individual subjects.

When comparing the Error of groups at the end of training session on *Post-Training* block, we found a marginal group effect (F_2,62_=2.6673, p=0.0774, η^2^_p_= 0.07923), with smaller errors for Group_-SRR_ [Group_-S_: 100.39±8.25, Group_-SR_: 109.56±7.89, Group_-SRR_: 83.45±8.25] (see Figure 2D). Note that this effect was significant in our previous study with a larger sample size (n=91; for more details see results section of Vassiliadis et al., 2021).

NEUROPHYSIOLOGY

### 3.2 End-training motor excitability changes

At the end of training, we obtained resting TMS measurements of MEP (mean amplitude and variability), SICI and UDP. First, we assessed changes in MEP amplitudes at the end of the training i.e., TMS_END_ (measured as MEP_MEAN_ and MEP_CV_ and expressed in percentage of TMS_PRE_) in the three groups of subjects. We noted that, at the end of training, all groups displayed MEP_MEAN_ amplitude values above 100% of pre-training (see Figure 3A)-Group_-SRR_: 114.15±10.83, Group_-SR_: 119.96±10.35, Group_-S_: 107.91±10.83.

**Figure 3.**
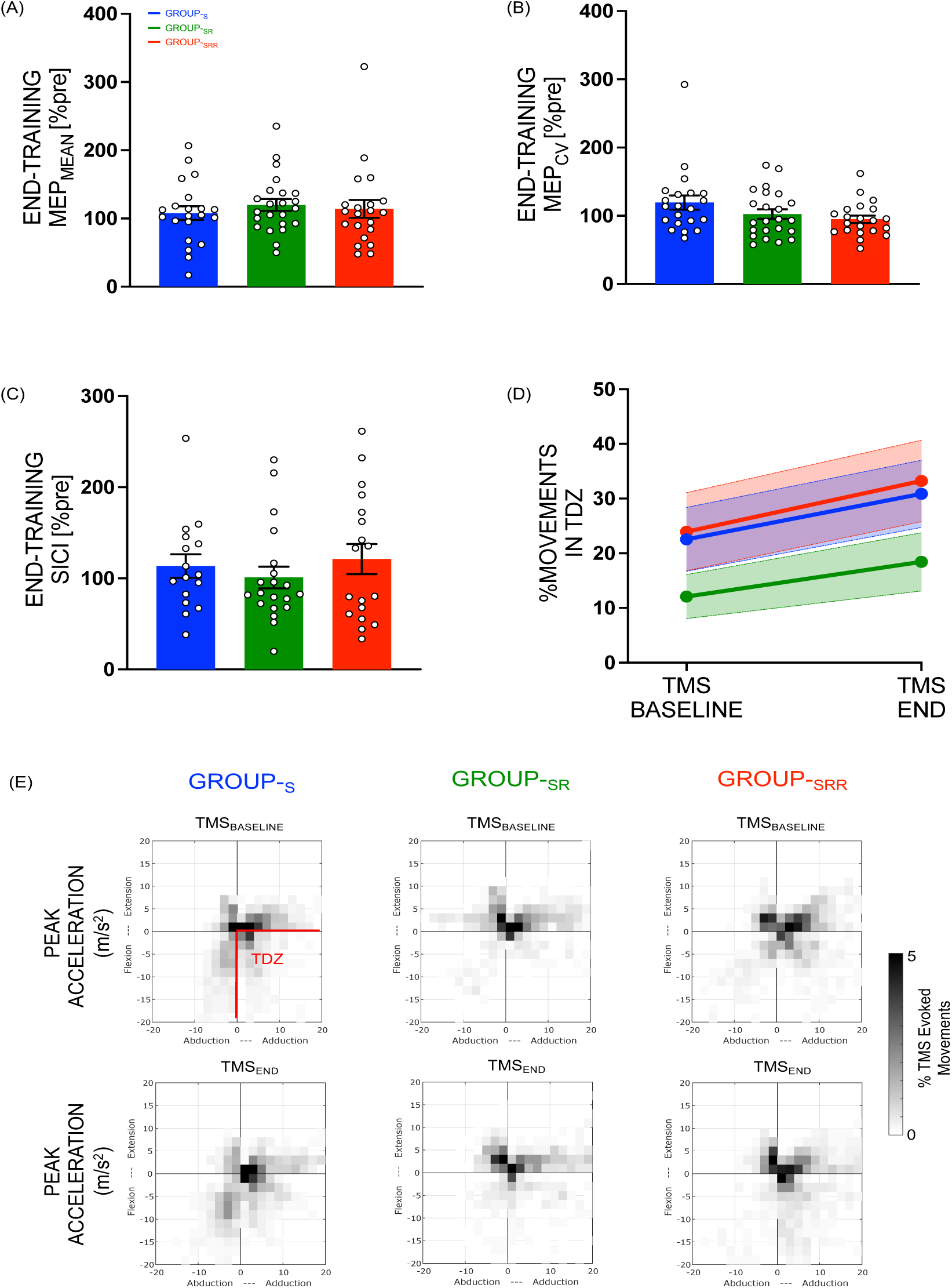
End-training motor plasticity measures: Resting TMS measurements obtained after Post-Training block when all three groups received same type of performance feedback on the skill task indicate no significant group differences for (A) MEP_MEAN_, (B) MEP_CV_, (C) SICI or (D) UDP. (E) Training led changes in UDP with more TMS evoked movements (measured using accelerometers in m/s^2^) obtained at TMS_END_ as compared to TMS_BASELINE_ in all groups in Training Direction Zone (TDZ) [refer to the grey scale on the right end for interpreting the number of movements in TDZ depicted in the grid plots). Bar plot data represent mean, normalized to TMS_PRE_. Error bars/bands represent Standard Error. White circles (with black outline) represent individual subjects.

However, one-way ANOVA did not reveal any significant effect of group on MEP_MEAN_ (F_2,62_=0.3232, p=0.7251, η^2^_p_= 0.0103). Independent of groups, we noted a significant effect of training on the MEP amplitude (single sample t-test against 100%: t_(64)_ = 2.3294, p=0.0230), suggesting training-induced increase in MEPs.

On the other hand, for MEP_CV_ at TMS_END_ time-point, we noted lowest variability of MEP in the group that received reward during training session [Group_-SRR_: 94.88±8.02, Group_-SR_: 102.48±7.66, Group_-S_: 119.44±8.02, see Figure 3B], with a non-significant trend (F_2,62_=2.4618, p=0.0936, η^2^_p_= 0.0735) revealed in the one-way ANOVA. Next, we measured SICI related changes in the three groups by computing the ratio of MEP_CONDITIONED_ and MEP_TEST_ pulses at the end of training i.e., TMS_END_ (expressed in percentage of TMS_PRE_). Statistically, we found no significant group effect (F_2,51_=0.5690, p=0.5697, η^2^_p_= 0.0218) [Figure 3C-Group_-SRR_: 121.13±13.88, Group_-SR_: 100.97±13.17, Group_-S_: 113.51±14.73]. We also pooled the groups together and performed a single sample t-test against 100% and did not find a significant change in SICI at the end of training (t_(53)_ = 1.4349, p=0.1572).

Finally, we assessed UDP and found that the percentage of movements in TDZ was higher at TMS_END_ time-point [27.50±3.47] as compared to TMS_BASELINE_ [19.52±3.47] (Figure 3D). At the group level, we noted that this value was numerically higher for Group-SRR [28.57±5.54], as compared to Group-SR [15.25±5.29] and Group-S [26.70±5.54]. Yet, upon statistical examination, our two-way ANOVA revealed a significant effect of time-point (F_1,62_=7.4016, p=0.0084, η^2^_p_= 0.107), but no significant group (F_2,62_=1.7959, p=0.1745, η^2^_p_= 0.055) or time-point X group interactions (F_2,62_=0.0885, p=0.9154, η^2^_p_= 0.003). This indicates that skill training led to UDP irrespective of the group type (see 3E for group-wise data of TMS evoked movements in TDZ).

In conclusion, at the end of training when all groups were back to performing the skill task with the same task performance feedback (no monetary reward and only reinforcement feedback) we did not find any reward related effects on MEPs, SICI or UDP for this time-point.

### 3.3 Early-training motor excitability changes

As mentioned earlier we were curious to know if exposure to reward influenced motor excitability measures (resting TMS measurements obtained for MEP mean and variability, and SICI) at this early-training time point which differs from the end of training when all groups performed the skill task with the same performance feedback. We, therefore, assessed MEP_MEAN_ and MEP_CV_ values at TMS_EARLY_ (expressed in percentage of TMS_PRE_), which fell at an earlier time point during the training session (between training block 2 and 3). Interestingly, at this early time point, there was a marginal group effect for MEP_MEAN_ amplitude (F_2,62_=2.6902, p=0.0758, η^2^_p_= 0.0798) with a trend for larger amplitudes at this early time point observed in the reward group [see Figure 4A-Group_-SRR_: 108.18±6.96, Group_-SR_: 85.90±6.65, Group_-S_: 97.98±6.96]. Moreover, for MEP_CV_ we noted lowest variability in the reward group-Group_-SRR_: 91.27±6.90, Group_-SR_: 101.13±6.59, Group_-S_: 117.95±6.90 (Figure 4B). Here, we found a significant effect of group (F_2,62_=3.8217, p=0.0272, η^2^_p_= 0.1097) and follow up post-hoc test revealed a significant difference between Group_-SRR_ and Group_-S_ (p=0.0220), while not between Group_-SRR_ and Group_-SR_ (p=0.5593) or between Group_-SR_ and Group_-S_ (p=0.1913). This effect of reward on MEP variability (when coupled with reinforcement and sensory feedback) particularly pronounced early during training could be consistent with its role in regulating motor variability, as previously shown behaviorally (Vassiliadis et al., 2021, Dhawale et al., 2017). Finally, early-training SICI did not differ between the three groups (Figure 4C; Group_-SRR_: 135.12±16.85, Group_-SR_: 120.28±15.98, Group_-S_: 108.70±17.87; no group effect: F_2,51_=0.5858, p=0.5604, η^2^_p_= 0.0224).

**Figure 4.**
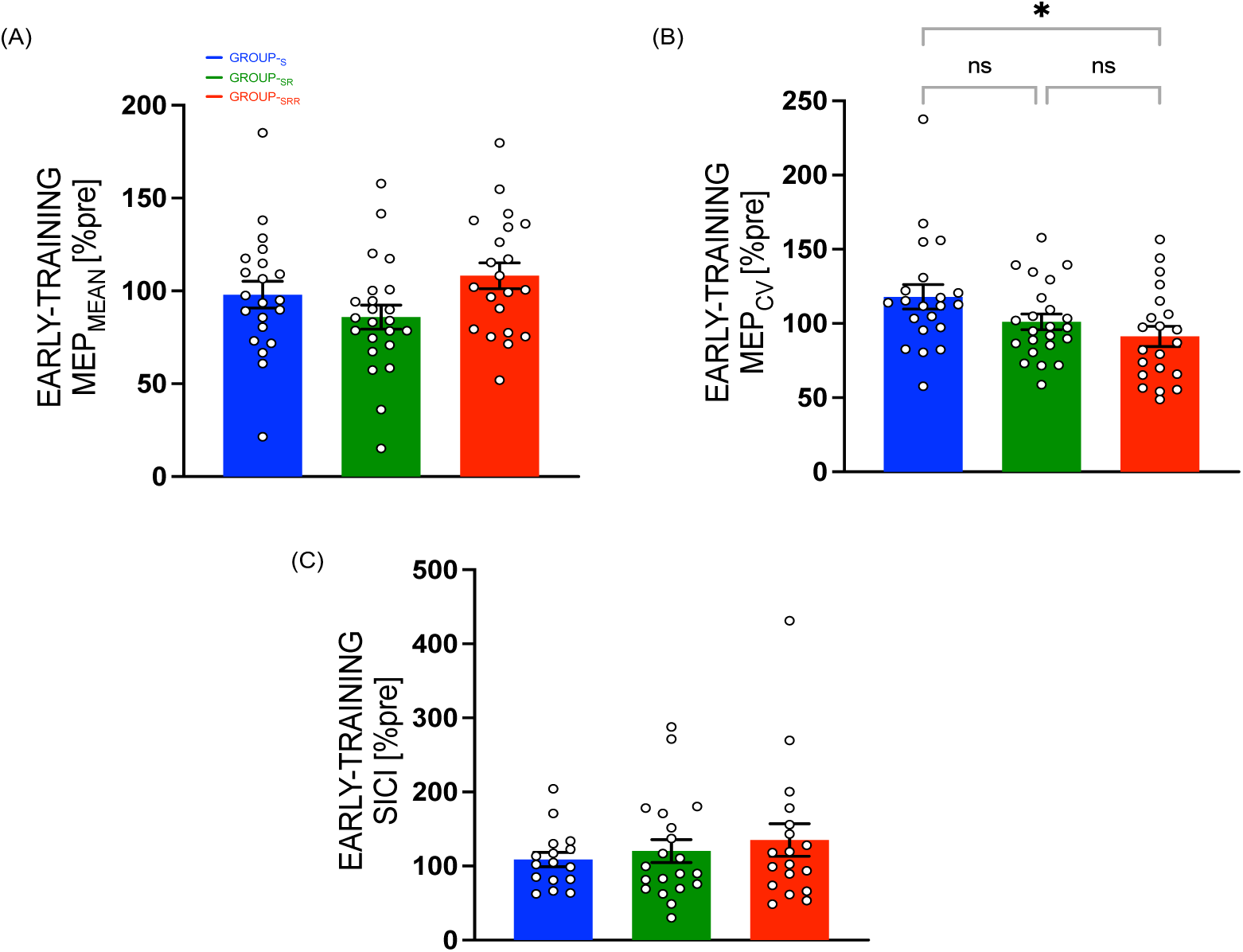
Early-training motor plasticity measures: Resting TMS measurements obtained early during Training Session when the three groups received different types of performance feedback on the skill task indicate (A) no significant group differences for MEP_MEAN_. (B) MEP_CV_ was significantly lower for Group-_SRR_. (C) no group differences observed in SICI. Bar plot data represent mean, normalized to TMS_PRE_. Error bars represent SE. White circles (with black outline) represent individual subjects.

In our protocol, TMS_END_ included more TMS pulses (i.e., 60) than TMS_EARLY_ (minimum 18, see methods) to allow us to reliably evaluate UDP when training has ended. Because we found a significant decrease in MEP variability only at TMS_EARLY_, in a control analysis, we asked whether this pattern of result could be explained by the different number of trials at both time points. To do so, we ran the same analyses on bootstrapped MEP values.

More specifically, we randomly selected without replacement 18 trials per time-point and calculated the mean MEP_MEAN_ and MEP_CV_ over 10000 resamples. Notably, even when using this approach, we found a similar pattern of results at TMS_END_ for MEP_MEAN_ (F_2,62_=0.3263, p=0.7229, η^2^_p_= 0.0104), with a comparable trend for a group effect in the MEP_CV_ data (F_2,62_=2.5512, p=0.0861, η^2^_p_= 0.0760). This control analysis indicates that the different number of trials per time point cannot explain our findings of lower MEP_CV_ at TMS_END_ Taken together, the data show an early-training reduction in corticospinal output variability (with no significant effect on mean CSE or GABAergic intra-cortical inhibition) when participants trained with reward.

### 3.4 Correlations between corticospinal excitability measure and sensitivity to reward-punishment scores in the reward group

In our above assessments on the role of reward on measures of motor plasticity, we found a significant effect of reward on the variability of MEPs i.e., MEP_CV_ at early-training TMS time point. We therefore wanted to further explore whether this MEP_CV_ obtained at TMS_EARLY_ in the reward group (GROUP_-SRR_) was associated with the individual sensitivity to reward and punishment measured obtained independently with the SPSRQ (see Methods). Upon performing bivariate Pearson correlation, we found a significant negative correlation between MEP_CV_ and sensitivity to reward scores (r(21)= −0.5231, p=0.0150). Note that this effect was not observed for Group_-SR_ (p=0.3361) or Group_-S_ (p=0.1917). On the other hand, for Group_-SRR_, no significant correlation was found between MEP_CV_ and sensitivity to punishment scores (r(21)= 0.0789, p=0.7337). This indicates that individuals with higher reward, but not punishment sensitivity scores in Group_-SRR_ had the largest reduction of MEP variability when measured during early training (Figure 5A and 5B).

**Figure 5.**
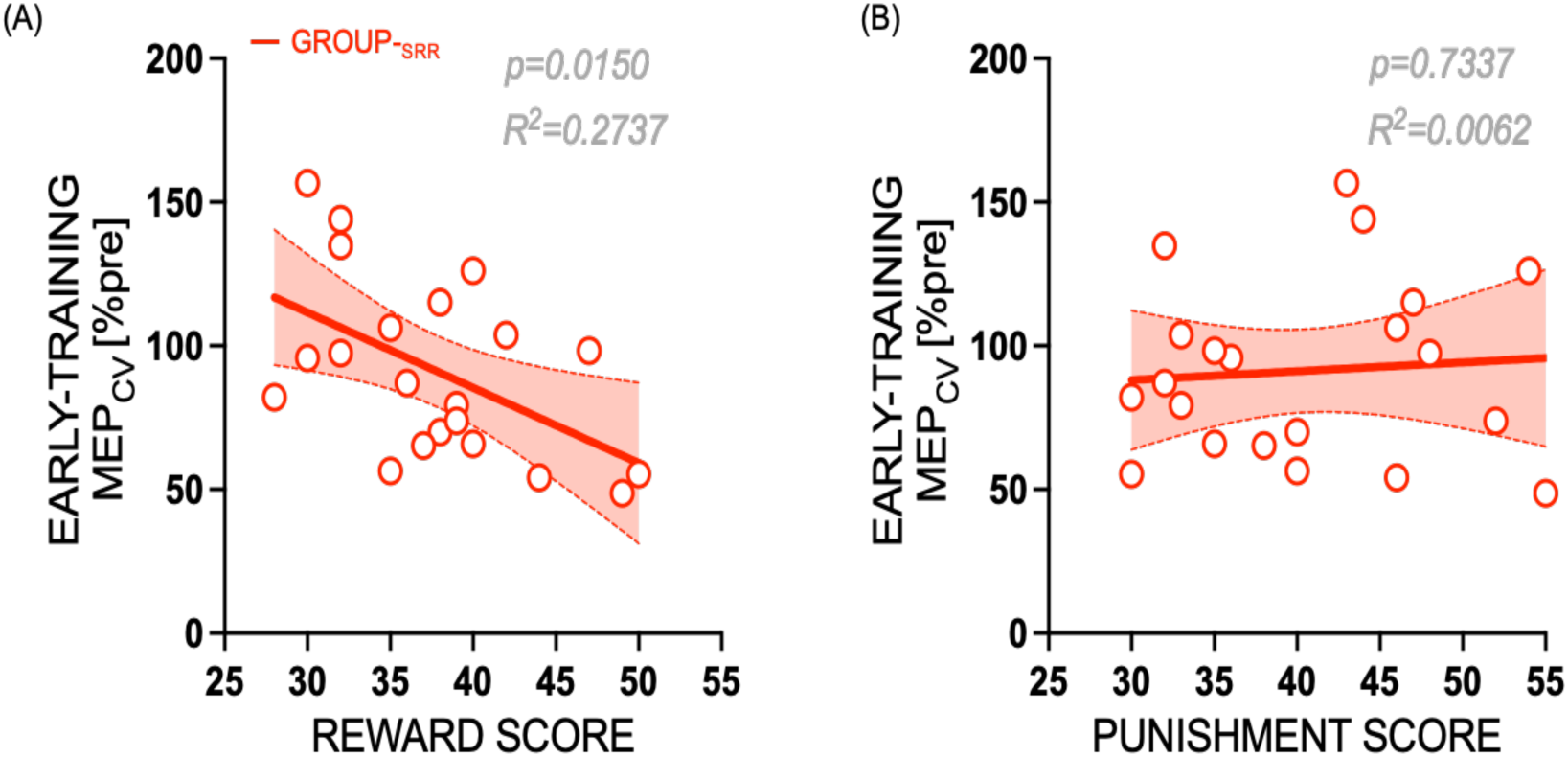
Early-training MEP_CV_ for individuals who received reward during the Training Session (Group-_SRR_) correlated significantly with (A) reward sensitivity scores (individuals with higher reward sensitivity score have lower MEP variability), but not with (B) punishment scores obtained using independent questionnaire (SPSRQ). Plot show linear regression fit (red solid line) with 95% confidence interval (red shaded band). White circles (with red outline) represent individual subjects of Group-_SRR_.

## 4. DISCUSSION

In this study, we set out to understand the motor neurophysiology underlying the benefits of reward on motor learning. In line with previous work (Vassiliadis et al., 2021; Sporn et al., 2022), we found that the presence of extrinsic reward during training accelerated learning compared to when training was performed with sensory feedback alone or coupled with reinforcement. Next, we assessed the effects of this extrinsic reward on motor plasticity measures by computing CSE (amplitude and variability of MEPs), SICI and UDP. While we did not find any significant change when evaluating motor plasticity after training, we found that subjects who learned the motor skill in the presence of extrinsic reward exhibited a reduction in CSE variability early on during training. Interestingly, this effect was correlated with individual subjects’ sensitivity to reward scores obtained on self-reported questionnaires. These findings suggest that faster skill learning in the presence of reward is driven by a reduction in the variability of corticospinal output and depends on individual personality traits.

The presence of extrinsic reward significantly reduced CSE variability early during training, and this effect remained at the trend level after learning. This result suggests that reward modulates a form of plasticity that is associated to more consistent resting-state CSE, ultimately facilitating skilled motor control. As such, neural variability in motor cortex in monkeys (Churchland et al. 2006a, 2006b) and CSE variability in humans (Klein-Flugge et al., 2013) decreases during action preparation and this reduction is associated to the efficiency of motor responses. Hence, the reduction of CSE variability we observe might reflect a refinement process that rapidly brings M1 firing rates closer to a specific optimal state for movement generation (Churchland et al. 2006a, 2006b; Klein-Flugge et al., 2013). In line with this idea, computational models suggest that the benefits of reward on motor control rely on a reduction of intrinsic neural noise that could increase the robustness of the corresponding motor representations (Manohar et al., 2015; 2019). Electrophysiological recordings during the task are required to better understand the effect of reward on neural noise during motor control and its relationship to the effect we report at rest.

Which neural processes could mediate this reward-dependent reduction of neural variability in the motor system? A possible mechanism may be plastic adjustments in circuits connecting hubs of reward processing such as the ventral tegmental area (VTA), the ventral striatum or the orbitofrontal cortex and M1 (Joel et al., 2002; McHaffie et al., 2006; Berridge, 2012). For instance, there is now evidence showing that M1 reward signals (Ramkumar 2016; Ramakrishnan et al., 2017; Levy et al., 2020) which are modulated during learning (Lee et al., 2022; Ghanayim et al., 2023) and causally involved in reinforcement motor learning (Levy et al., 2020), originate, at least in part, from a dopaminergic pathway linking VTA and M1 (Hosp et al., 2011; Leemburg et al., 2018; Ghanayim et al., 2023). Importantly, VTA-M1 reward signaling seems to be particularly important in the early stages of motor skill acquisition in rodents, but not when a plateau of performance is reached (Hosp et al., 2011, Leemburg et al., 2018), in line with the learning stage-dependent effect of M1 disruption during reward-based decision-making in humans (Derosiere et al., 2017a, 2017b). Consistently, a recent rodent study found that the VTA-M1 pathway was causally involved in the reorganization of M1 in the early stages of a reinforcement motor learning, allowing M1 activity to evolve rapidly towards an expert configuration (Ghanayim et al., 2023). Hence, an interpretation of our result is that the early reduction of corticospinal variability may be related to the more consistent inputs reaching M1 when training with reward. Interestingly, this effect also scaled with the individual’s sensitivity to reward. This is consistent with previous observations that sensitivity to reward (as indexed by the SPSRQ) may reflect inter-individual variability in the structure (see Barros-Loscertales et al., 2006) and function (Adrian-Ventura 2019) of key hubs of the reward network (e.g., VTA, ventral striatum, orbitofrontal cortex), possibly modulating their influence on M1 activity during reinforcement skill learning. More research is required to better understand inter-individual factors shaping behavioral and neural responsiveness to reward during motor learning, an aspect that could be promising to determine which patients could benefit from reward-based motor rehabilitation protocols. Overall, our data suggest that the early reduction of neural variability in motor cortex may be an important mechanism underlying the benefits of reward on motor learning.

Although we did find a significant increase of CSE amplitude irrespective of the group, in line with previous work (e.g., Pascual-Leone et al. 1995; Butefisch et al. 2000; Duque et al. 2008; Galea and Celnik 2009; Christiansen et al. 2018; Vassiliadis et al., 2020), this training-related modulation was not influenced by reward. This result may seem to contrast with previous literature showing reward-related modulations of CSE amplitude in reaction time tasks (Gupta and Aron, 2011; Klein et al., 2012; Freeman et al, 2014; Bundt et al., 2019). Yet, an important difference here is that in these studies, CSE was evaluated during motor preparation within the task, while we measured plasticity at rest, between blocks of training. Hence, a possibility is that reward-related modulations of CSE amplitude during a task do not translate to a persistent change in CSE, in line with a previous study on a smaller sample (Mawase et al., 2017). In this scenario, changes in resting-state CSE amplitude and variability may reflect the operation of distinct mechanisms, possibly reflecting the modulation or either the firing rates of neurons or their consistency during rest (Churchland et al., 2006a). Our results suggest that reward-based motor learning preferentially modulates the resting-state variability of CSE rather than its amplitude.

In addition to CSE, we investigated the putative effect of reward on SICI and UDP. Indeed, to explore plastic changes reflecting intracortical interactions, we assessed whether GABAergic SICI can be modulated by reward in our task. We observed no reward-driven effect either early during training or at the end of training on SICI. A recent work by Hamel et al (2023) shows that monetary reward decreases SICI during movement preparation in a motor sequence task but interestingly such effect is not observed in their control experiment in which SICI is measured post-movement when the reward feedback has already been processed. This may explain the lack of effect on SICI in our data, where our SICI measurements were made at rest after the training session was over. Our findings imply that intracortical interactions in the presence of extrinsic reward may diminish at rest. Therefore, studies interested in these measurements and reward-based motor skill learning should be designed accordingly. As for UDP, which was measured at the end of training, we noted a practice-induced change (relative to baseline), consistent with prior studies showing plastic changes based on repetition of specific movements toward a target direction (Classen 1998; Duque et al., 2008; Diedrichsen et al., 2010; Huang et al., 2011; Bernardi et al., 2015; Mawase 2017). However, we did not find any group difference related to reward. One plausible explanation is that all three groups in our study repeated the same movement and comparable UDP at the end reflects repetition-based, as opposed to success-based, Hebbian changes in M1 (Bütefisch et al., 2000; Orban de Xivry et al., 2011; Verstynen and Sabes, 2011; see Exp-1 of Mawase et al. 2017). Our results thus do not show an interplay between UDP and extrinsic reward-based reinforcement learning related to M1 for the skill acquired in our study.

Finally, we would like to address some factors that could have influence our results. First, our study was not optimized to isolate subtle changes in UDP. Unlike previous studies on UDP, our force modulation task involved isometric, and not ballistic, movements and the training direction was not opposite to the TMS-induced movement direction, as classically done (Classen et al., 1998; Duque et al., 2008). Still, our study supports the view that classical skill force modulation tasks can elicit UDP (Mawase et al., 2017). Second, in this study only the group that trained with reward and reinforcement (Group_-SRR_ as compared to Group_-SR_ and Group_-S_ who trained without reward) exhibited significant skill learning, thus the effects of reward on plasticity measures reported here should be interpreted in the context of learning. Taken together, we report that the presence of reward during skill training induces an early reduction of corticospinal variability at rest during motor skill learning and that this effect depends on individual’s sensitivity to reward. This study supports the view that motivation by reward boosts specific plasticity mechanisms in motor cortex during motor skill learning and adds to the growing body of literature showing reward-related activity in M1 (e.g., Ramakrishnan et al., 2017; Levy et al., 2020). In a broader perspective, these results fit well in the framework of embodied cognition (Foglia and Wilson, 2013; Sullivan, 2018) in which motor behavior and the associated neural mechanisms can be influenced by cognitive processes such as motivation, urgency, emotions or pain.

## Acknowledgments

We would like to thank Aegryan Lete and Wanda Materne for assistance with data acquisition.

